# GPSA2: combining landmark-free and landmark-based methods in geometric morphometrics

**DOI:** 10.1101/2024.08.03.604701

**Authors:** Benjamin J. Pomidor, Matt Dean

## Abstract

Geometric Morphometrics (GM) revolutionized the way that biologists quantify shape variation among individuals, populations, and species. Traditional GM methods are based on homologous landmarks that can be reliably identified across all specimens in a sample. However, landmark-based studies are limited by the intensive labor required of anatomical experts, and regions of interest are often devoid of landmarks. These limitations inspired the development of many “landmark-free” approaches, but unreliable homology estimation and complicated underlying mathematical bases can make biological interpretation challenging. Here we present GPSA2, a novel method for analyzing surface meshes that combines landmark-based and landmark-free methodology within the familiar framework of Generalized Procrustes Analysis. In a major innovation, our method can incorporate user-defined landmarks into otherwise landmark-free analysis by transforming the landmarks into pointwise shape descriptors that are exploited during iterative homology estimation and superimposition (i.e. “alignment” of objects). GPSA2 also addresses a longstanding issue in morphometrics – the impact of variability in the distribution of sampled points over an object – by introducing a surface area-weighted shape distance metric and superimposition cost function. The improved homology approximation, together with the application of Taubin smoothing and an optional resistant-fit superimposition technique, ensure robust analysis even when a dataset exhibits regions of intense shape variation. We apply GPSA2 to two empirical datasets: 15 primate skulls and 369 mouse bacula. Our analyses show that inclusion of landmarks increases biological accuracy, and that GPSA2 produces summaries of shape variation that are easy to visualize and interpret.

## INTRODUCTION

In the decades since its inception, the field of Geometric Morphometrics has provided the fundamental mechanisms for describing and interpreting variation in size, shape, and form between individuals, populations, and species [1–7]. Traditionally, the raw data for Geometric Morphometrics comes in the form of biologically homologous landmarks that are reliably identified, then digitized as 2D or 3D coordinates from every object in a sample (e.g., the infraorbital foramina across a set of skulls). Procrustes superimposition [8] computationally aligns two objects by minimizing the combined distance between corresponding landmarks. The remaining difference represents shape.

Unfortunately, landmark-based approaches require significant labor from an anatomical expert, bottlenecking the acquisition of data. Furthermore, identifiable landmarks are not equally distributed on an object, and regions devoid of landmarks drop out from any downstream analyses even though they might contain important shape variation [1]. Even when an object features ample landmarks, subjective decisions about which landmarks to include for a specific research objective must be considered.

Relatively recently, several “landmark-free” methods have been developed that alleviate some of these limitations [9–17]. These methods perform superimposition and downstream analyses on the entire digital representations of objects, typically in the form of point clouds or surface meshes made of thousands to millions of points, rather than the dozens of individual landmarks typical of a traditional geometric morphometric dataset. Landmark-free approaches include shape information across the entire object and do not require manual digitization of homologous points. This comes at a cost, however: these methods rely on computational approximations of the true biological homology, and even small errors in this approximation can have severely detrimental impacts on morphometric analysis [for detail, see Section 11 of 1]. In some cases, the complex mathematics that underlie landmark-free approaches muddle biological interpretation [1], which is otherwise straightforward in standard landmark-based geometric morphometrics.

Here we present a novel hybrid approach that combines landmark-free concepts with landmark-based methodology to establish a landmark-informed, pointwise homology approximation for a set of surface meshes. The key advance is the addition of local shape descriptors, which are computed values that describe aspects of shape around a point on a mesh – for example local curvature [18]. Local shape descriptors themselves are not new, but the novel inclusion of local shape descriptors in a Procrustes-like framework offers a clearly interpretable mathematical path to incorporate user-defined landmarks as local shape descriptors in an otherwise landmark-free method. Our approach builds on multiple foundational concepts laid out in our previously published landmark-free method, GPSA [15]. This new method, which we dub GPSA2, integrates several significant advances over GPSA, including the incorporation of local shape descriptors and landmarks, the derivation of a new shape distance metric & superimposition cost function to account for uneven distribution of points on a surface mesh, and the application of a smoothing algorithm to reduce distortion in the computed mean surface during Procrustes superimposition. We also developed a novel space-partitioning tree to improve computational speed in searches for best-matching points, as well as an alternative superimposition technique for resistant-fit in the case of large-scale non-homologous features (e.g., a horn present on some skulls but not others).

GPSA2 combines landmark-based and landmark-free techniques within the well-established and easily interpretable mathematical framework of Generalized Procrustes Analysis. After describing the algorithmic development in detail, we demonstrate GPSA2 with two datasets: the 15 primate skulls originally analyzed in Pomidor et al. [15] and the 369 mouse bacula analyzed by Schultz et al. [19]. Analysis of these datasets reveals the advantage of our hybrid approach, and shows that GPSA2 yields clear, robust descriptions of shape variation.

## MATERIALS & METHODS

### Overview of GPA, GPSA, and GPSA2

In traditional geometric morphometrics, reliably present, repeatably identifiable points called *landmarks* are digitized by an anatomical expert, defining a pointwise *homology map* from one object to another (e.g., the intersection of two sutures on one skull is assumed to be homologous to that same intersection on another skull) (Fig. 1) [20]. Typically, *Procrustes superimposition* is performed to computationally overlap two objects by minimizing the root sum of squared distances between each pair of homologous landmarks. The remaining root sum of the squared distances between corresponding landmarks is called *Procrustes distance*, and is used to quantify the difference in shape between the objects. *Generalized Procrustes Analysis* (GPA) extends this comparison from two objects to a set of objects [21]. By superimposing all the objects onto a *reference shape* (an artificial object made of the landmark averages across the dataset), GPA avoids the O(*n*^2^) superimposition problem that would arise if all pairwise alignments were evaluated. Once superimposition is complete, the Procrustes distance between any pair of objects can be readily calculated, and the variation in the location of landmarks across the dataset is the *shape variation*.

**Fig. 1.**
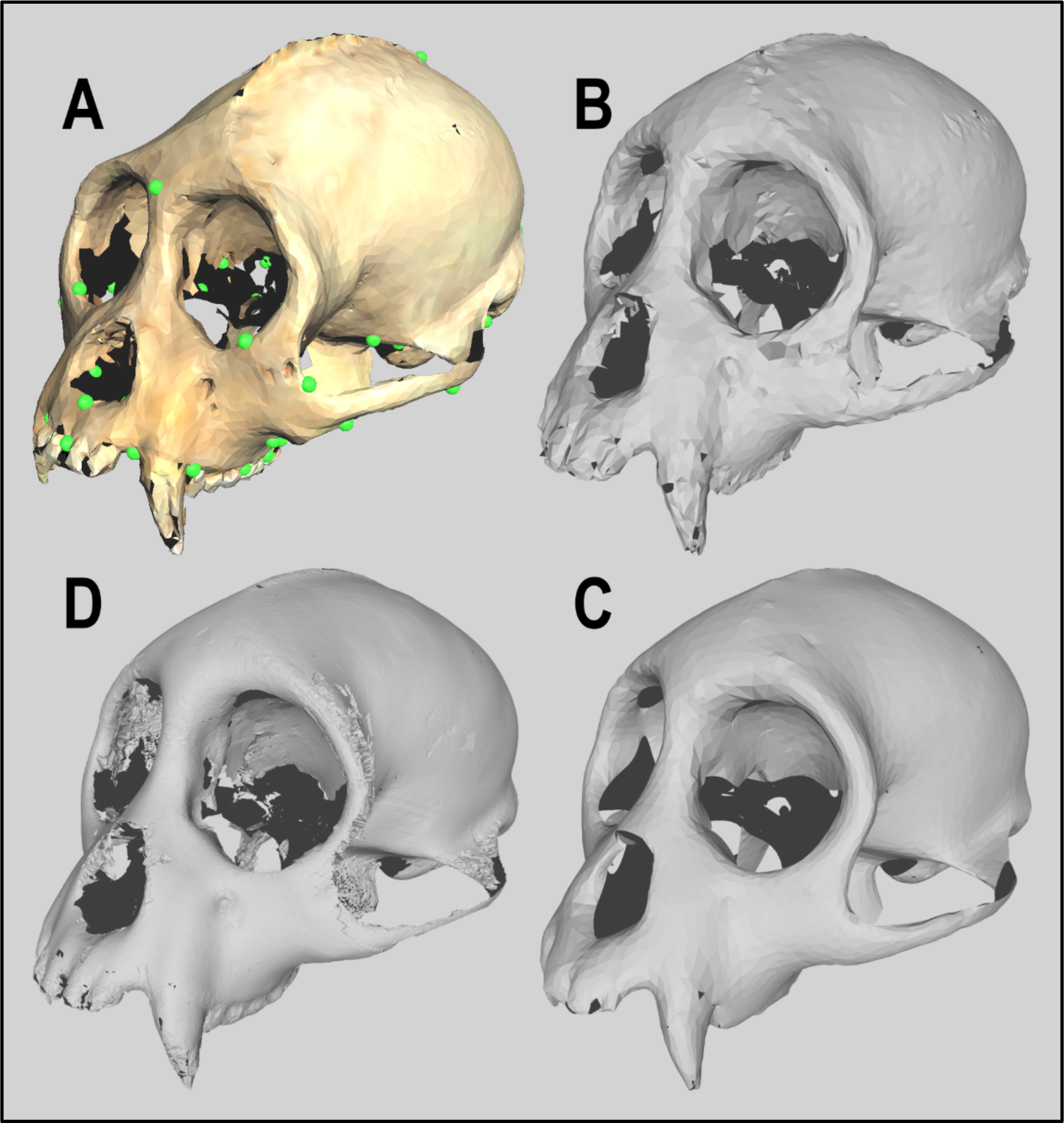
Various primate skull surface meshes. (A) Locations of the 43 landmarks represented by green circles on the skull used as the reference surface. (B) The reference surface after superimposition and deformation to fit the mean sample shape, but prior to the application of Taubin smoothing. (C) The reference surface after superimposition, deformation to fit the mean, and Taubin smoothing has been applied (Kpb = 0.01, N = 50). (D) The mean reference surface extracted by GPSA in Pomidor et al. [15] exhibiting instability in regions of high variation.

*Generalized Procrustes Surface Analysis* (GPSA) [15] is an entirely landmark-free morphometric analysis method that operates on surfaces made of *point clouds* (collections of unmeshed 3D points) rather than landmarks. Without landmarks, superimposition is more challenging because the homology map between two objects is undefined. GPSA solves this via a modified symmetric version of the Iterative Closest Point (ICP) algorithm, iteratively alternating between approximating the homology map between two objects and superimposing them, until the superimposition has converged to a local minimum [22,23]. GPSA extends this pairwise superimposition process to a sample size greater than two in a similar manner to GPA — by superimposing each object in the sample (sets of landmarks for GPA, point clouds for GPSA) onto a reference object, then recomputing the reference to fit the mean shape of the dataset. The resultant averaged point cloud is then used to define an approximate homology map across the entire dataset.

The new method we present here, dubbed *GPSA2*, operates on surface meshes instead of point clouds. Meshes include edges that connect adjacent points on a surface, providing a more complete representation of an object and facilitating the use of local geometry to improve approximated homology maps. Like GPSA, GPSA2 applies a two-phase iterative process alternating between a pairwise homology map approximation step and a superimposition step. In the approximation step, GPSA2 uses a combination of the position of a point on a mesh, the vector normal to the surface at that point, and any number of user-defined *local shape descriptors* to more accurately search for the best-matching point on the opposing mesh. Local shape descriptors can be computed values based on geometry near a point (such as curvature) [24–26] or user-defined landmarks converted to mesh-wide descriptive variables, enabling hybrid landmark/surface mesh analysis, a key capability of GPSA2.

In the superimposition step, GPSA2 deploys a novel shape distance metric and superimposition cost function that accounts for the surface area of the faces connected to each point, eliminating sampling density bias that could arise with large variance in face size across a mesh. Taubin smoothing [27] applied during the reference object deformation portion of generalized superimposition helps prevent noisy distortion when fitting the reference surface to the sample mean. Finally, we incorporate an auxiliary superimposition method for resistant-fitting based on resistant-fit [28–30] and Trimmed ICP [31] for datasets with large, non-homologous features, such as a horn that is present on some, but not all, skulls in a sample. These non-homologous features introduce extra shape variation that may be incorrectly “spread” over the rest of the object under standard Procrustes-style superimposition [32]. Resistant-fit techniques significantly reduce this effect by removing the contribution of non-homologous regions to superimposition.

We discuss each of these advancements in more detail below, then analyze two different datasets to illustrate the capabilities of GPSA2. GPSA2 is freely available on GitHub (https://github.com/bjpomidor/GPSA2) as a configurable command-line program with scalable parallel processing, under Apache License 2.0, along with both datasets presented here, under the Creative Commons Attribution 4.0 International license. Our GPSA2 library was written in Java 7 using Apache Commons Math 3.6.1 [33].

### Phase 1: Homology map approximation

The goal of this phase is to map approximately homologous points to each other across a set of surfaces in 3D space. For every point on each surface object, we identify the best-matching point on a designated reference object (alternately referred to in the literature as an “atlas” object). We also do the reverse: for every point on the reference object, we identify the best-matching point on each surface object in the sample. This matching procedure is done in both directions to ensure the relationship between any two objects is symmetrical, a necessity for measuring and comparing shape distances. This collection of point-pairings between two surfaces defines a homology map. By calculating the homology maps between each object in the sample and a single reference object, we can extrapolate the map between any pair of objects from the sample [34,35].

The criteria for best-matching points varies widely from method to method. In landmark-based Procrustes analysis, it is unnecessary to approximate the homology map because the landmarks are already homologous by definition, so the best match is already known. In the ICP formulation underlying GPSA, using the closest point in 3D space is sufficient for convergence [22]. In GPSA2, where the goal is to map objects of varying shape to one another for comparison, we use local shape descriptors to improve the estimation of the best-matching point. The term “local shape descriptor” encompasses a wide variety of techniques [18], but at its core refers to values that describe the shape of an object near a given point. For example, a typical surface mesh consists of triangular faces connected by their common edges and vertices; from just this information we can calculate a normal vector for each triangular face [36]. If the normal vectors for adjacent faces are parallel, then we know that region is relatively flat. If they diverge, we know the region is curved. Measuring this divergence gives us a numerical value for curvature [24], which can be used to compare shape similarity between points.

Under the assumption that homologous points occur in regions with relatively similar shape, incorporating local shape descriptors can improve homology approximation. GPSA2 allows the user to define how many and what type of local shape descriptors to use and weight them preferentially to fit their dataset. A huge diversity of local shape descriptors has been proposed in the literature [37–39], but a review by Heider et al. [18] concluded that a descriptor based on the distribution of normal vectors in combination with a descriptor based on curvature metrics generally produced the best performance, which is the strategy we adopt below.

Local shape descriptors provide a framework to incorporate landmark data into otherwise landmark-free analysis. In GPSA2, each landmark is converted into a separate local shape descriptor, the value of which is calculated at each point on the surface mesh. Our library uses geodesic distance from the landmark for this conversion, but further exploration of other landmark-to-descriptor conversion functions is warranted.

To build a homology map between two surfaces, GPSA2 evaluates the similarity between pairs of points with local shape descriptors via a standardized similarity measure *D_S_* (related to Mahalanobis distance [40]). For points *a* and *b* on two different objects with coordinates *x*, *y*, and *z* in Euclidean space, *n_x_*, *n_y_*, and *n_z_* in normal space, *l_0_* … *l_d_* in local shape descriptor space and weighting terms *w_n_* (for normal vectors) and *w_l_0* … *w_l_d* (for local shape descriptors):

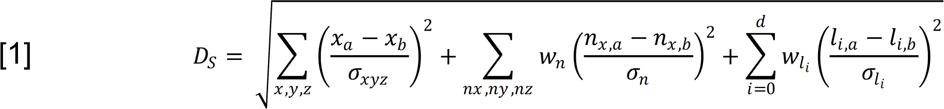

The three summation terms inside the square root correspond to the contribution of Euclidean distance between the 3D coordinates of *a* and *b*, the differences in their normal vectors, and the differences in their local shape descriptors, respectively. Each term is normalized by its standard deviation (σ_xyz_, σ_n_, and σ_li_), as they are otherwise not directly comparable; one unit of Euclidean distance is not equivalent to one unit of curvature. Evaluating *D_S_* for every possible pair of points to determine which pairings are the best match would be computationally expensive (O(*n^2^*)), so we developed a novel space partitioning tree to reduce search complexity to O(*n*·log *n*) [41] (see A1 for detail). Once the best-matching vertex is found, the surface of the connected faces are checked for a closer interpolated match to negate issues stemming from large faces and varying sampling density. For example, a point that is close to the center of a large triangular face will likely have a better match at its projection on the surface of the triangle than at any of the distant vertices.

Together, the local shape descriptors, similarity measure *D_S_*, and the space partitioning tree provide efficient, effective means to approximate the homology maps required for the second phase of the algorithm.

### Phase 2: Superimposition

The objective of this phase is to computationally superimpose all the objects in the sample as closely as possible, relying on the homology maps generated in the previous phase. A pair of surface objects and the homology map between them are fed into a superimposition cost function that, when solved, gives us a set of transformation parameters to optimally align one surface object with another. Superimposing all the objects in the sample on a single reference object is almost as good as superimposing them all on each other pairwise, but computationally much cheaper [21]. The relative positioning of the sample objects after such a superimposition heavily depends on the reference object, but we can reduce this bias by deforming the reference object to fit the mean shape of the sample and repeating the superimposition process. Typically, a few repetitions of this process are sufficient to produce a stable mean shape, but selection of the initial reference object must be done cautiously. Objects with regions that are missing or that exhibit significantly different shape from the rest of the sample can see their specific differences propagate through the generalized superimposition process and bias the result [42].

Once the sample is superimposed, we can measure the shape distance between two objects using a mesh-based distance metric. For two surface meshes *A* and *B*, made up of *m_A_* and *m_B_* points, with total surface areas *a_A_* and *a_B_*, where *p_A,i_* is the *i*th point on *A* with an associated surface area weight of *w_A,i_* and *q_B,i_* is the corresponding homologous point on *B*, and where *p_B,j_* is the *j*th point on *B* with an associated surface area weight of *w_B,j_* and *q_A,j_* is the corresponding homologous point on *A*, the shape difference between *A* and *B* is defined by the equation:

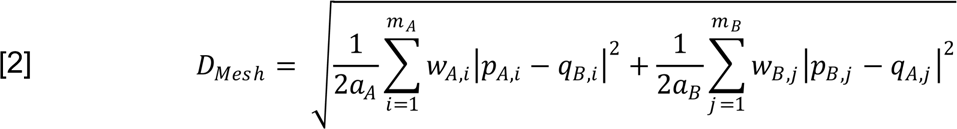

Equation 2 is essentially the Procrustes surface metric from GPSA with a key modification: The addition of a pointwise weighting term determined by the surface area of each point’s connected faces ensures that variance in sampling density over an object does not bias GPSA2 to favor more densely sampled regions, a potential issue for some landmark-free techniques [43]. This equation retains the reciprocal structure of the Procrustes surface metric, incorporating distances calculated using both the homology map from *A* → *B* as well as from *B* → *A*, because as with GPSA, the homology maps are neither reciprocal nor exclusive: a single point *b* on surface *B* could be the best match for multiple different points on *A*, while an entirely different point on *A* might be the best match for point *b*. Including both maps is a simple method to ensure the metric is symmetric and usable for downstream analysis. Note that the weighting terms here are entirely separate from the weighting terms used in similarity measure *D_S_*; distance metric *D_Mesh_* is based solely on Euclidean distance and surface area.

We derive our rigid superimposition cost function directly from this metric as a function of rotation and translation to ensure that the shape distance is minimized when superimposition is complete. Size is removed beforehand (as much as possible) by scaling by the inverse of an area-weighted size estimate *s_a_*:

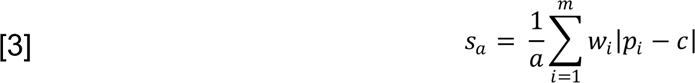

Where *a* is the surface area of the object, *m* is the number of points, *w_i_* is the surface area for the *i*th point on the surface, *p_i_*, and *c* is the area-weighted centroid of the object, similarly calculated. The initial value of *s_a_* is recorded for each object for use in downstream analysis. With size addressed, the distance metric is converted into a cost function by dropping the now-extraneous square root and applying a rotation matrix *H* and translation vector *t* to one surface, *B*:

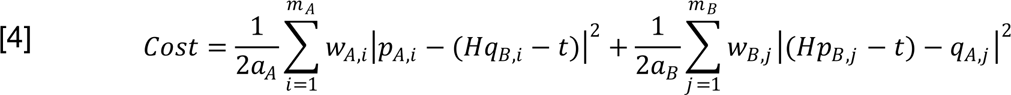

We minimize this cost function by linearizing *H* in terms of rotations about the x, y, and z axes (*r_x_, r_y_,* and *r_z_,* respectively) and building a system of equations from the partial derivatives with respect to each transformation parameter, set to zero. We are left with:

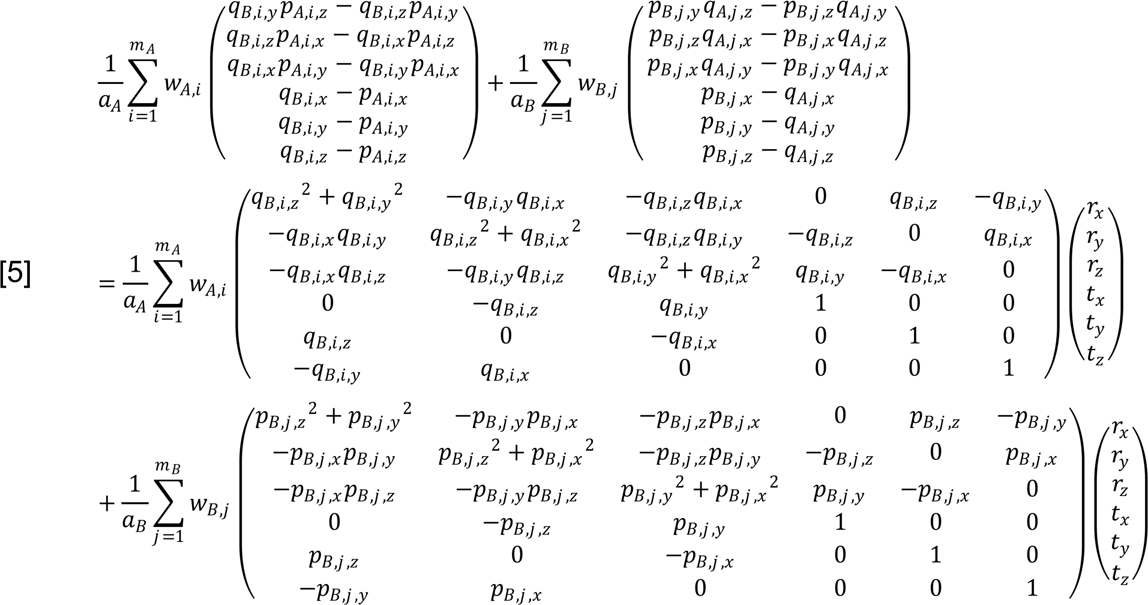

After calculating the coefficients, we solve the 6×6 linear system and apply the resulting transformations to surface B, bringing the two surfaces closer to superimposition. As with the iterative closest point algorithm, the approximate homology maps (Phase I) and superimposition transformation parameters (Phase II) are alternately re-calculated and reapplied, iteratively, until the superimposition has converged.

Once all surfaces have been jointly superimposed on the reference object, we use the homology maps to calculate the average surface for the sample. Not only is this useful for downstream analysis and visualization methods, but we use this new average surface to refine our superimposition by replacing our reference object with it for another round of superimposition in a similar manner to GPA. Unfortunately, the discretized nature of the surface mesh makes this calculated average prone to having discontinuities and a “noisy” surface texture, with the severity dependent on both the smoothness of the homology maps and the quantity/quality of the sample surface data. To counter this, we apply Taubin smoothing [27], a parameterized mesh geometry signal processing technique that effectively filters the high-frequency shape “noise” from the averaged surface (Fig. 1). Combined with the better homology map approximation offered by the addition of local shape descriptors, the stability gained from this smoothing step significantly improves the quality of the average surface generated by GPSA2 over the original point-cloud-based GPSA (Fig. 1).

Especially large local shape variation or non-homologous shape data in a sample (ex: horns present on some skulls but not others) presents a statistical difficulty: the shape-metric-minimizing superimposition described above effectively averages such shape variation over the rest of the object to possible detriment [44]. To analyze this type of sample, we developed an alternate superimposition method based on Slice’s resistant-fit method for landmarks [30] and the Trimmed Iterative Closest Point algorithm (TrICP) [31]. This new resistant-fit superimposition method identifies and leaves out the significantly different regions of two objects, evaluating the cost function only for the regions relatively conserved on both surfaces. We do this by sorting the best-matches in the homology map between the two objects by an estimated *match quality* value that describes how similar a matched pair of points are to each other and their neighbors (see A2 for detail). We then use the golden section search algorithm [45] to identify the appropriate cutoff to maximize the proportion of the objects used for superimposition while minimizing poorly matched points; matches that fall outside of this cutoff are excluded from the superimposition cost function. The end result is a superimposition that constrains the excess shape variation from high-variation/non-uniform features to the features themselves, rather than spreading it over the rest of the object. There are other challenging geometric morphometric issues that most current landmark-free methods currently cannot resolve [46], but GPSA2’s basis in the generalized Procrustes framework may prove similarly advantageous in adapting existing landmark-based solutions to those problems in the future.

After superimposition is complete, the results are transformed for use in further statistical analysis. The superimposition is summarized via an interspecimen distance matrix containing the shape distance (described above) calculated for every pair of individuals in the sample. The approximated homology map is converted into usable statistical data by deforming the final reference surface into a series of *homologized surfaces* that each exactly fit one of the individuals in the sample. The vertices of these surfaces, the *homologized points*, are used to construct an *n* x 3*m* (*n* individuals, *m* points in three dimensions on the reference surface) matrix akin to a matrix of shape variables from a landmark dataset, though care should be taken with interpretation in the face of such “high-density” data [47]. We can perform PCA on this homologized points matrix to obtain ordinations that efficiently summarize the shape variation in the sample, while projection along each PC axis results in a surface that coherently illustrates the regional distribution of the shape variation.

### Example dataset: Primate skulls

This dataset was included in the original publication of GPSA [15]. Here, we evaluate GPSA2 using the same n=15 male *Cebus apella* skulls, which have both surface scans and landmarks (m = 43). We tested our method without descriptors to examine the impact of the new area-weighted point-to-mesh correspondence, then with basic local shape descriptors (normal vector, Gaussian & mean curvature, and curvature index [25]), and finally with landmark-derived local shape descriptors to determine the effect on homology map approximation.

The mean surface, the homologized surfaces, a variance heatmap, and projected surfaces for the first two principal axes were generated for the landmark-descriptor test run and visualized using Meshlab [48] to subjectively assess the results. The difference in superimposition and shape distance measurement was quantified in two ways: first, by measuring the displacement of each landmark from its location in a GPA superimposition when superimposed with the surface-based methods (GPSA, GPSA2 with no descriptors, GPSA2 with simple local shape descriptors, GPSA2 with local shape descriptors and landmark-based descriptors), and second, by calculating the elementwise correlation of interspecimen matrices for Procrustes distance (landmarks), GPSA’s surface metric, and GPSA2’s area-based surface metric. Both sets of comparisons were visualized in R [49] with the original GPSA results included as a baseline.

### Example dataset: Mouse bacula

Schultz et al. [19] performed a quantitative genetics study on size and shape variation of 369 bacula (penis bones) in a panel of 75 recombinant inbred lines of mice. Each baculum was represented as a cloud of points with 3D coordinates, segmented from microCT scans. GPSA2 takes surface meshes as input, so we wrote a small program to convert the point clouds back to voxel data (VoxelConverter.jar, https://github.com/bjpomidor/GPSA2), then “shrinkwrapped” these voxel data to extract a surface mesh.

The size of each surface mesh was calculated via the area-weighted size metric described above. To measure variation in shape, we first standardized all surface meshes by this size measure, then completed superimposition and homology approximation using the GPSA2 software. We first analyzed the data in a purely landmark-free way, then repeated the analysis after including a single landmark to examine the utility of even a limited application of the landmark/local shape descriptor framework. This landmark was manually digitized as the “peak” of a protrusion that occurs at the distal and dorsal position of the baculum (Fig. 2). This landmark was the only one that we could reliably identify from all specimens. We analyzed the homologized points in a Principal Components Analysis (PCA), using the PRCOMP function in R, with center=TRUE and scale=FALSE and subsequently treated Principal Components as quantitative traits describing shape.

**Fig. 2.**
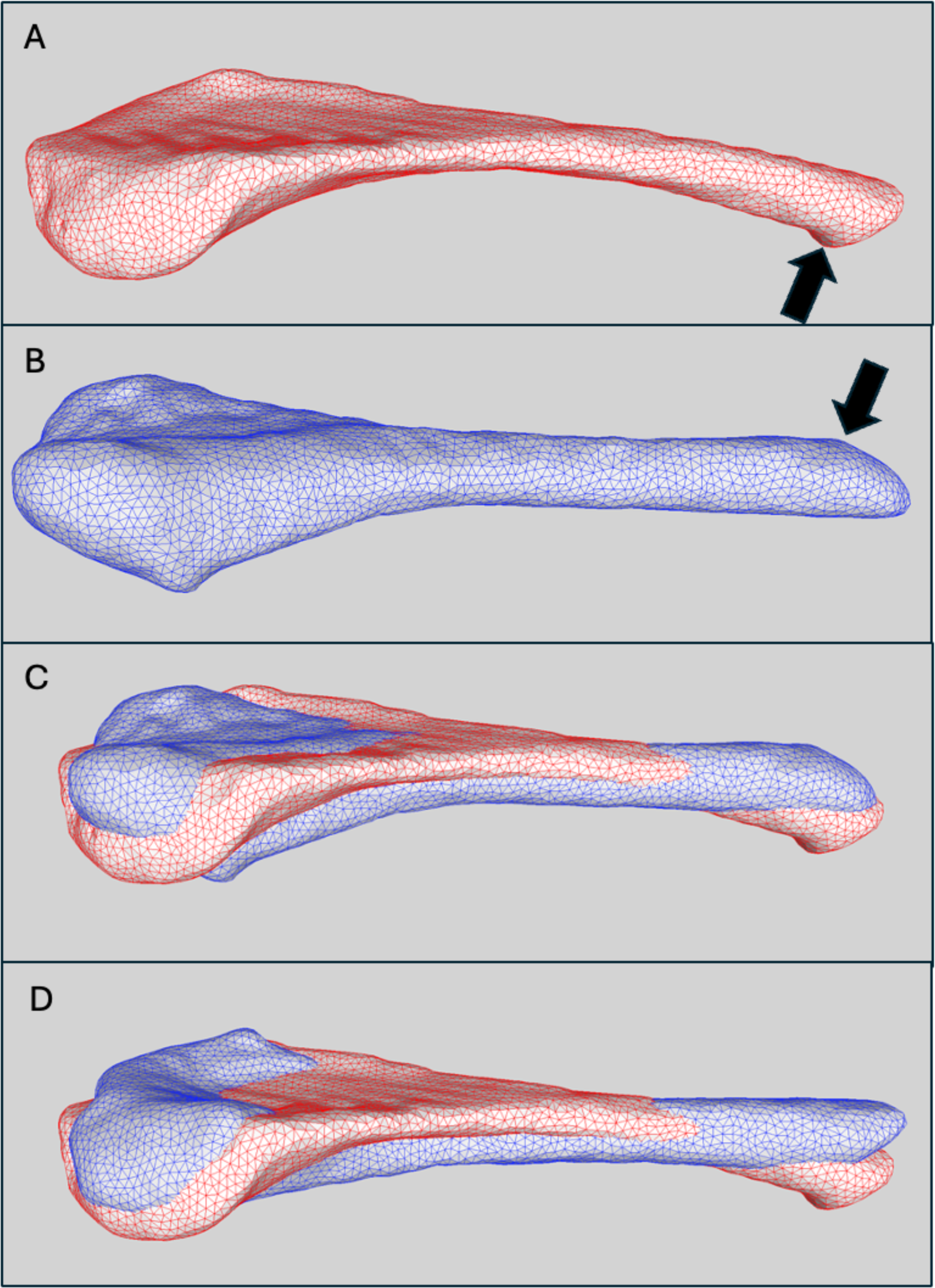
Surface meshes of mouse bacula illustrate several key concepts. All images are lateral views. In a purely landmark-free approach, the best-fit superimposition results in a “flipped orientation”: specimen C57BL_6J_012814.32.PT3000.IP (A) and specimen BXD21_02APRIL2015.22.PT2000.IP (B) align in a dorsoventral orientation (C). After incorporating a single landmark (black arrow), these two specimens align “correctly” (D), meaning that although the overall alignment is costlier, the two specimens are now oriented in the same direction.

Our original study [19] identified quantitative trait loci influencing size and shape variation. We repeated that analysis here but using GPSA2 output. Using either the area-weighted size values or the Principal Components as shape features, we employed the SCANONE function in the R package QTL, using Hayley-Knott regression to estimate the location and the effect of QTL [50]. To determine significance, we permuted phenotypes and genotypes 1000 times, and took the 95th quantile of the 1000 maximum LOD scores as our empirical significance threshold.

## RESULTS

### Primate skulls

GPSA2 outputs a variance map that is useful in visualizing the distribution of variance across the aligned objects. Much of the variation is located towards the interior of the skull (Fig. 3), likely stemming from occlusion artifacts when the skulls were originally scanned. Despite this, the surfaces generated by projection of the homologized points matrix PC axes display significantly more relevant shape variation in the zygomatic arch, frontal bone, and facial region (Fig. 3), a result with some precedent [51]. In this dataset, aggressive Taubin smoothing (Kpb = 0.01, N = 50) was required to effectively filter out shape “noise” (resulting from imperfect homology approximation) in the reference deformation and surface homologization steps (Fig. 1) and ensure stability with the aforementioned scanning artifacts. This resulted in some loss of detail, notably around the teeth.

**Fig. 3.**
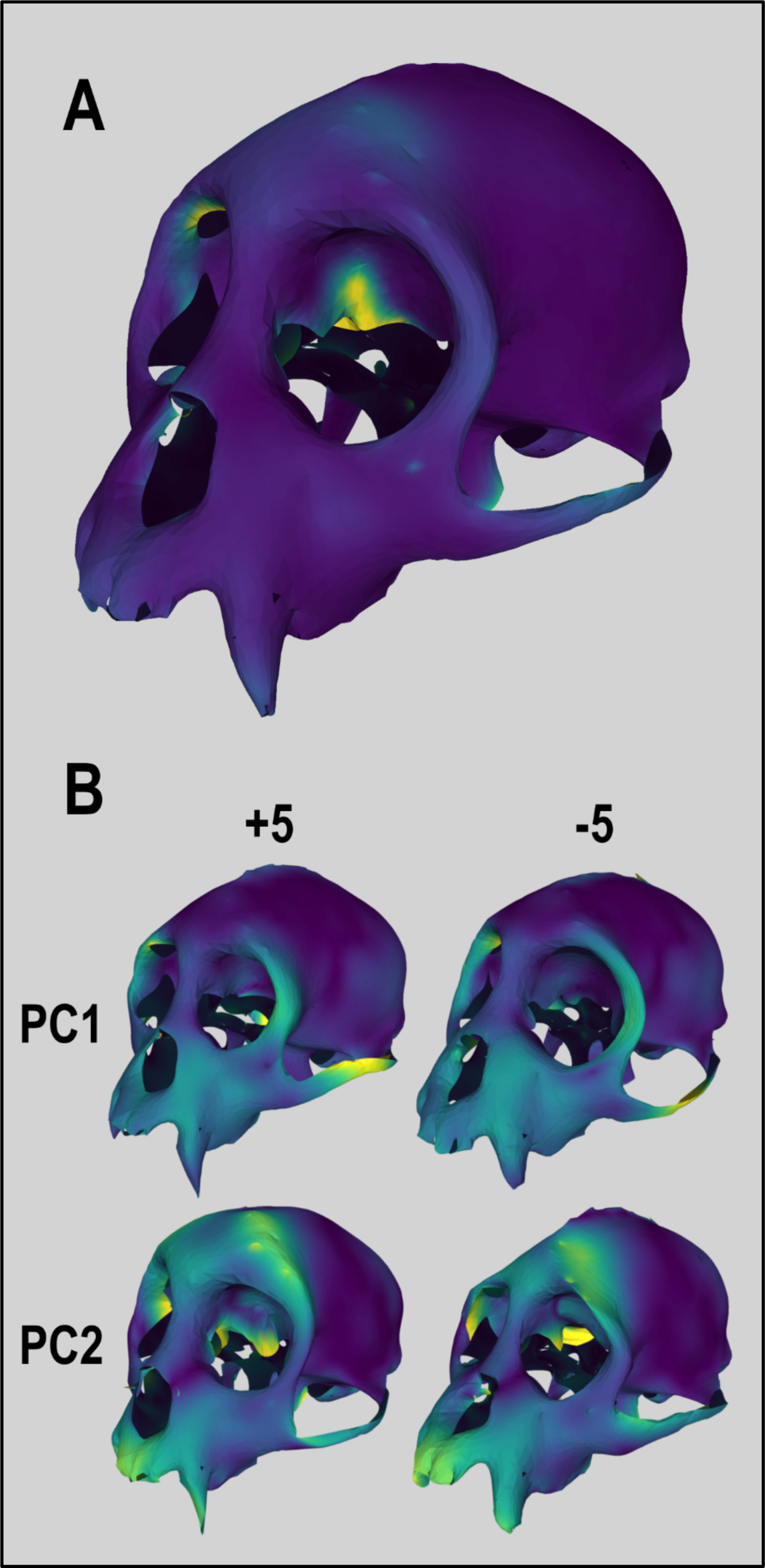
Variation and ordination heat maps visualized on primate skull surface meshes. (A) : The total shape variation for the sample, constrained largely to the interior structure. (B) : The first two PC axes (0.145, 0.135) projected at +5.0 and −5.0.

The distribution of landmark displacements (Fig. 4) indicate that both the transition from point clouds in GPSA to surface meshes in GPSA2 as well as the addition of local shape descriptors and landmark-based descriptors resulted in configurations slightly closer to the GPA superimposition. While the median values fall within a small range, there is still a noticeable (∼15%) difference between the median displacement distance for GPSA (0.00248) and GPSA2 with landmark-descriptors (0.00210). The elementwise interspecimen distance correlation (Pearson) plots (Fig. 5), which were all evaluated against the Procrustes distance matrix from the GPA superimposition, show marginal difference between GPSA and GPSA2 (with landmark-descriptors) with Procrustes distance matrices (R = 0.95 and 0.96, respectively). This implies that the GPSA and GPSA2 superimpositions are not dramatically different from GPA, nor from each other, at least in this dataset. However, when examining correlation with GPA based on each method’s respective surface-based shape distance, there was a noteworthy increase in correlation (R = 0.30 for GPSA, R = 0.41 for GPSA2). Both correlation values are expectedly weak because the surfaces represent very different shape data from the landmarks (Fig. 1); all but a few landmarks are located in the facial or ventral region. From these results we can infer that the improvements to GPSA2 facilitated better homology map approximation than in GPSA, underscoring the benefit of incorporating local shape descriptors and landmarks into an otherwise landmark-free approach.

**Fig. 4.**
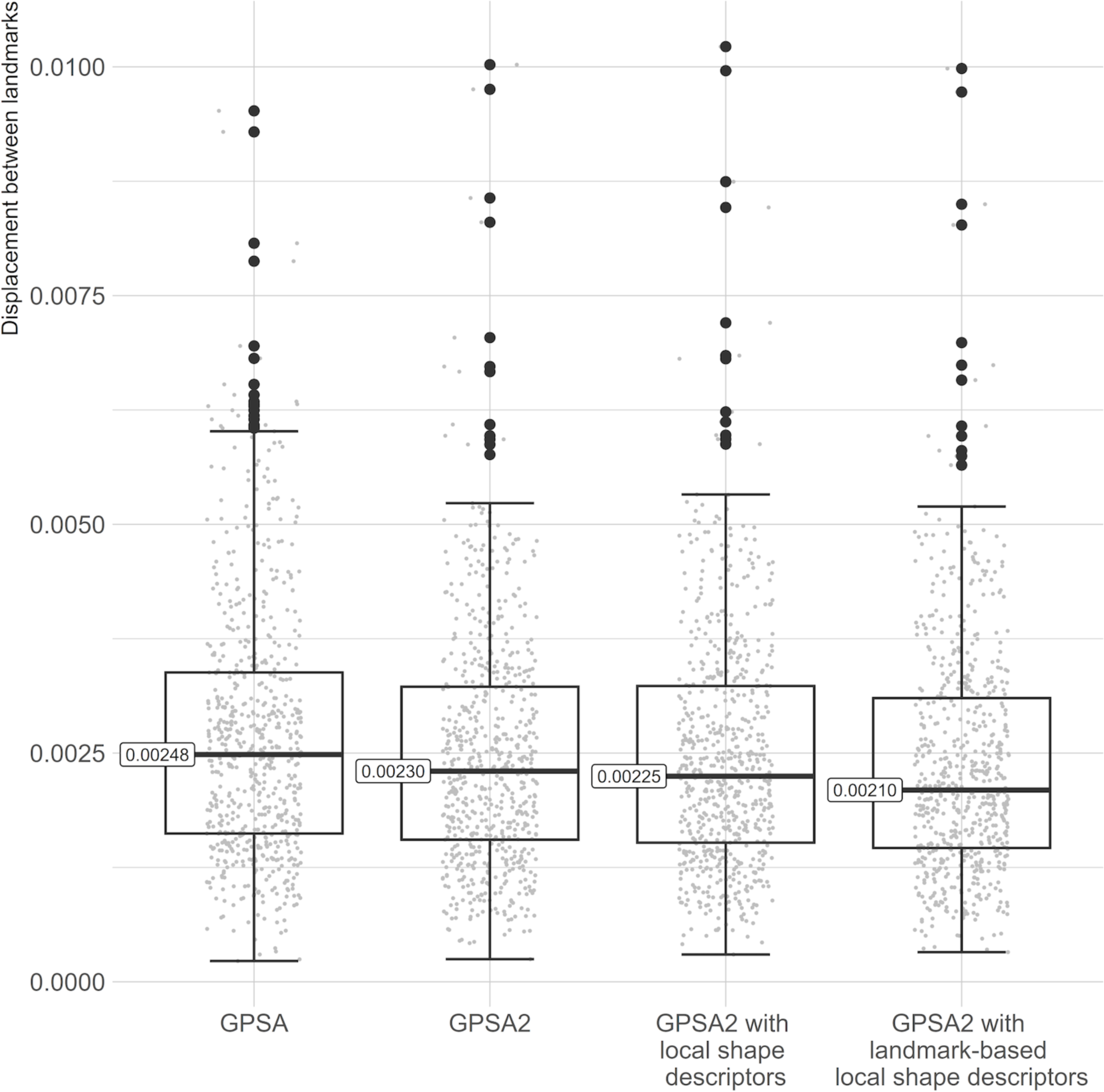
Box plot of per-landmark displacement by method vs GPA. The difference in position between all landmarks on all individuals superimposed with GPA and the same landmarks on the same individuals superimposed with other methods when the consensus landmark configurations are aligned. The lower the value, the closer that particular landmark on that particular individual is to its position in the ground truth GPA superimposition.

**Fig. 5.**
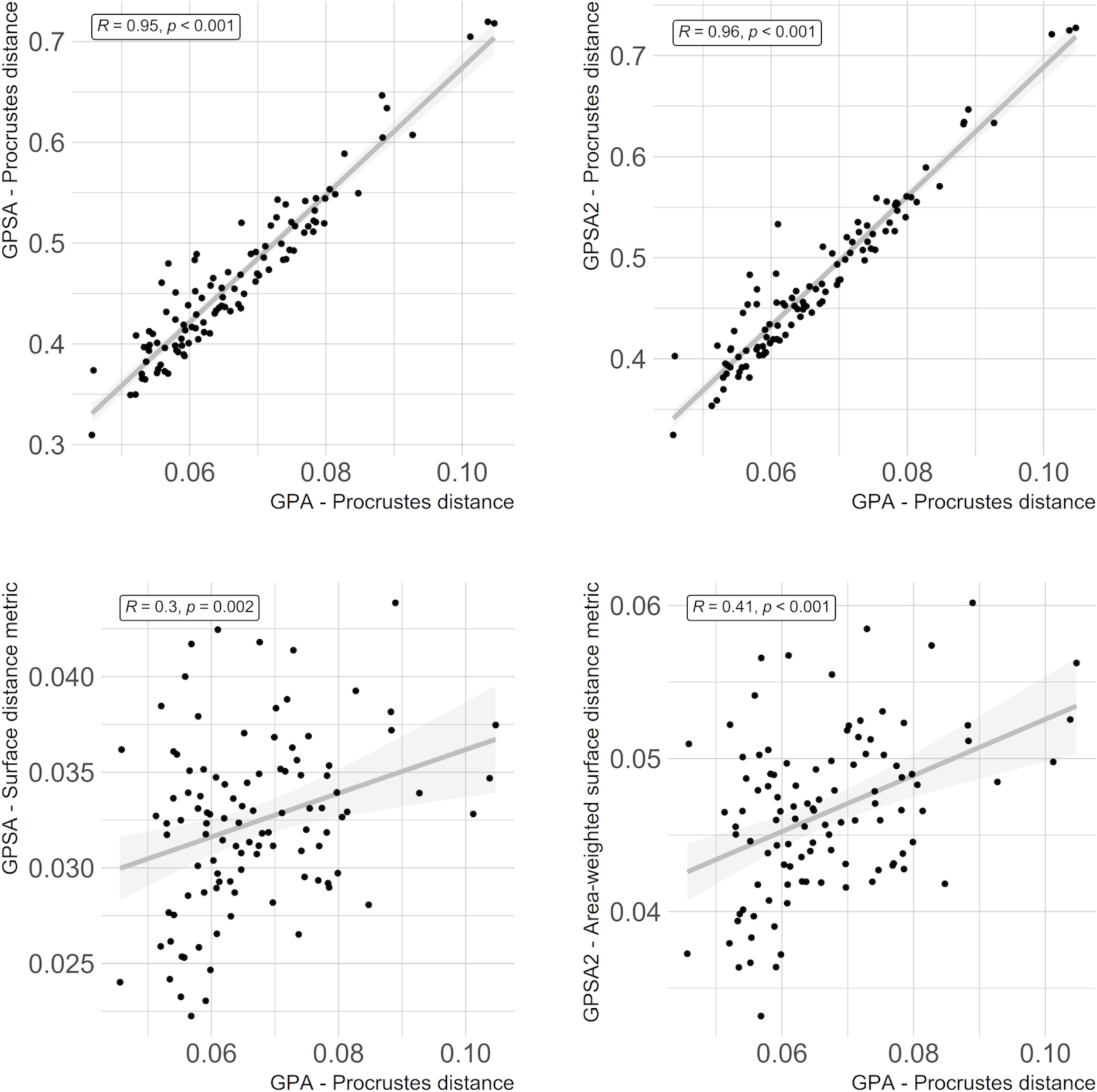
Various interspecimen distance matrices vs the GPA-superimposed Procrustes distance matrix, elementwise. Top row: The Procrustes distance matrices for both GPSA (left) and GPSA2 (right) superimpositions are highly correlated elementwise with the Procrustes distance matrix for a GPA superimposition (Pearson; R=0.95 and 0.96, respectively). Bottom row: The surface-based distance matrix from GPSA (left) is weakly correlated elementwise with the Procrustes distance matrix for a GPA superimposition (R=0.3). For the area-weighted surface-based distance matrix from GPSA2 (right), this correlation is stronger (R=0.41).

### Mouse bacula

When we ran GPSA2 as a purely landmark-free approach, the ventral side of one baculum sometimes aligned to the dorsal side of another (Fig. 2). This is likely due to the complex and variable curvatures of different bacula that sometimes resulted in the flipped alignment yielding a lower cost function even though it violated any reasonable definition of homology. Inclusion of the single digitized landmark fixed this issue (Fig. 2), demonstrating the advantage of combining landmark-free with landmark-based approaches.

The overall variance map shows that most of the variation across samples localizes to the distal tip as well as the lateral flanks of the proximal bulge of the baculum (Fig. 6). PC1 explained 43.4% of the variance in shape, ranging from relatively narrow to relatively wide proximal ends (Supplementary File 1, ventral view) as well as relatively straight to relatively bent (Supplementary File 1, lateral view). No significant QTL explained variance in PC1, suggesting that the variance in PC1 was not genetically determined. PC2 explained 16.4% of the variance in shape, mostly concentrated in the midshaft of the baculum (Supplementary File 1). Two significant QTL were associated with PC2, one on chromosome 2 and one on chromosome 8 (Supplementary File 2). PC3 explained 10.5% of the variance in shape, mostly concentrated in the “valley” that is characteristic of the dorsal, proximal side (Supplementary File 1). One significant QTL on chromosome 1 was associated with PC3 (Supplementary File 2).

**Fig. 6.**
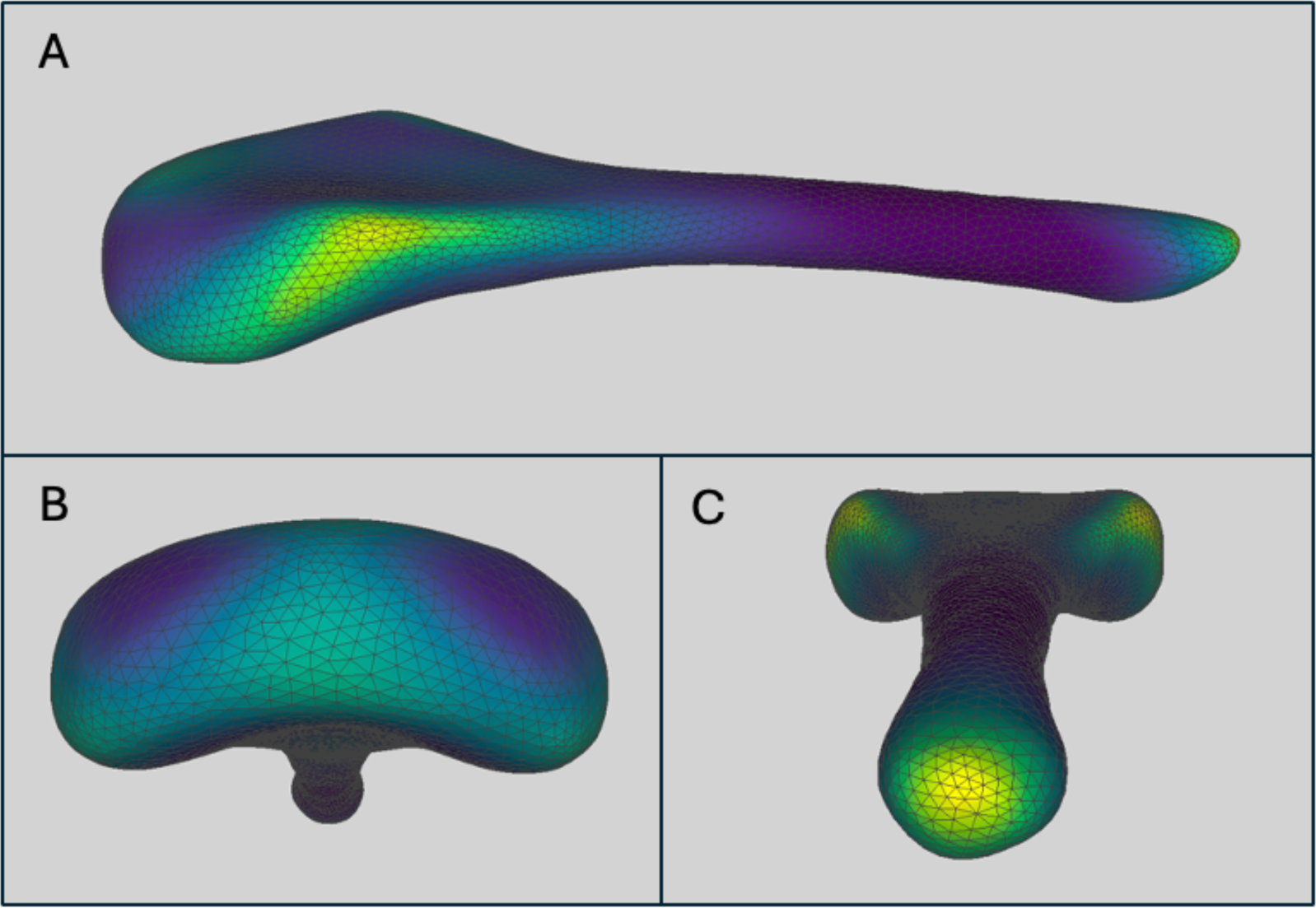
Lateral (A), proximal (B), and distal (C) views of the overall variance map produced as standard output from GPSA2. The baculum represents the average shape across all 369 specimens. Regions of high variance (yellow) are concentrated along the lateral edges of the proximal end as well as the distal end. See Supplementary Figure 1 for variance maps stratified by Principal Components.

## DISCUSSION

### Primate skulls

In our original publication of GPSA, analyses of primate skulls had one particularly undesirable result. Because the exact orientation and morphology of the zygomatic arch varied so much across the dataset, GPSA achieved little reciprocity in the approximated homology map. As a result, when averaged, the zygomatic arch “collapsed” and became unrealistically thin (Fig. 1), which then propagated across the set of homologized skull surfaces. While the zygomatic arch was by far the worst occurrence, this issue with nearest-neighbor-based homology was repeated to a lesser degree in other regions of high variation. GPSA2 fixed this problem with the addition of local shape descriptors, landmark-based descriptors, and mesh-based Taubin smoothing, all of which improve homology approximation and stability. However, these improvements required careful tuning of the Taubin smoothing parameters to preserve as much detail as possible while maintaining mesh stability.

The additional information captured by surfaces compared to landmarks is another matter for consideration. In this dataset, there are relatively few landmarks located on the dorsal side of the cranium; they are instead located mostly on the ventral side or in the facial region. This means we can reasonably expect a landmark-free method to do a better job at analyzing the shape of the un-landmarked portion of the cranium. Despite producing similar superimpositions, the disparity in shape representation between landmarks and surfaces accordingly resulted in weak correlation between landmark-based and surface-based shape distances. GPSA2 notably achieved slightly better alignment and moderately better distance matrix correlation than GPSA, implying that the improvements targeting homology map approximation were successful. This dataset conveniently demonstrates the downside to this all-encompassing shape analysis as well: the variance map (Fig. 3) showed large amounts of variation on the interior of the skull, where occlusions in the original scans made a jagged mess of little analytical value and largely independent of most of the rest of the shape of the object. Dimensional reduction via PCA pushes the variation from these artifacts to be combined with the more morphologically relevant variation, and, from the ordination projections (Fig. 3), do not appear to dominate any of the first several axes. Further investigation into how GPSA2 affects various types of error would be appropriate, in line with recent geometric morphometric-specific interest in measurement error [52].

Finally, the primate skull dataset needed to be decimated prior to analysis in order to achieve runtimes reasonable for development (<1 hour), which unfortunately resulted in some loss of detail due to the decreased resolution. Because each local shape descriptor adds another dimension to the data matrix for a given surface object, the jump in dimensionality from basic 3-dimensional coordinates to 3D coordinates plus normal vectors, measures of curvature, and 43 additional landmark descriptors was very large. With adequate processing power, memory, and time, though, there is no reason the method or software could not handle larger, higher-resolution datasets. In the future, since most of the surface is far away from most of the landmarks in a typical dataset resulting in values of zero or nearly zero, sparse matrix methods may be implemented to help reduce the extra computational load generated by additional landmark descriptors.

### Mouse bacula

Re-analysis of the baculum dataset succinctly illustrated the benefit of combining landmark-free and landmark-based approaches. The addition of a single digitized landmark completely corrected the occasional dorso-ventral flipping that happened with purely landmark-free methods.

Although a thorough quantitative genetics treatment is outside the scope of the current manuscript, our re-analysis demonstrates another major advantage of GPSA2. Because the homologized points can be analyzed in a Principal Components framework, orthogonal shape variance can be analyzed as separate phenotypes. In our original study [19] we uncovered a single significant QTL on chromosome 2 associated with PC2 shape variation. We identified this same locus associated with PC2 generated from GPSA2 analyses. However, we discovered two additional QTL using GPSA2: one on chromosome 1 associated with PC3 and the marginally significant one on chromosome 8 associated with PC2. Our original analysis combined shape variation into a single measurement; because GPSA2 enables Principal Components Analysis to separate shape variance into orthogonal axes, complex phenotypes may “interfere” less with each other’s signal, potentially providing more discriminatory power to QTL analysis.

### Conclusions

GPSA2 represents an intermediate step between the previous generation of geometric morphometrics and the developmental frontier of a new one. Manually placed landmarks are reliable and analysis via Generalized Procrustes Analysis is readily interpretable. Yet, the new generation of easily available, high-resolution surface and volumetric scans demand powerful landmark-free methods that rely on automated homology approximation predicated on complex mathematical models. Such methods are powerful, but may obfuscate the path to biological interpretation. Here GPSA2 finds its niche: it is a hybrid approach capable of extending landmark data into a surface-based context and works within a Procrustes-style framework that maintains ease of interpretation and downstream analysis. As described and demonstrated here, it achieves this through a novel combination of robust iterative superimposition and descriptor-based homology approximation.

## Acknowledgements

We thank Peter Beerli for his encouragement and guidance. Dennis Slice provided critical insights into the development and application of these methods. This work was funded by National Science Foundation grant #2027373

---

**Supplementary File 1.**
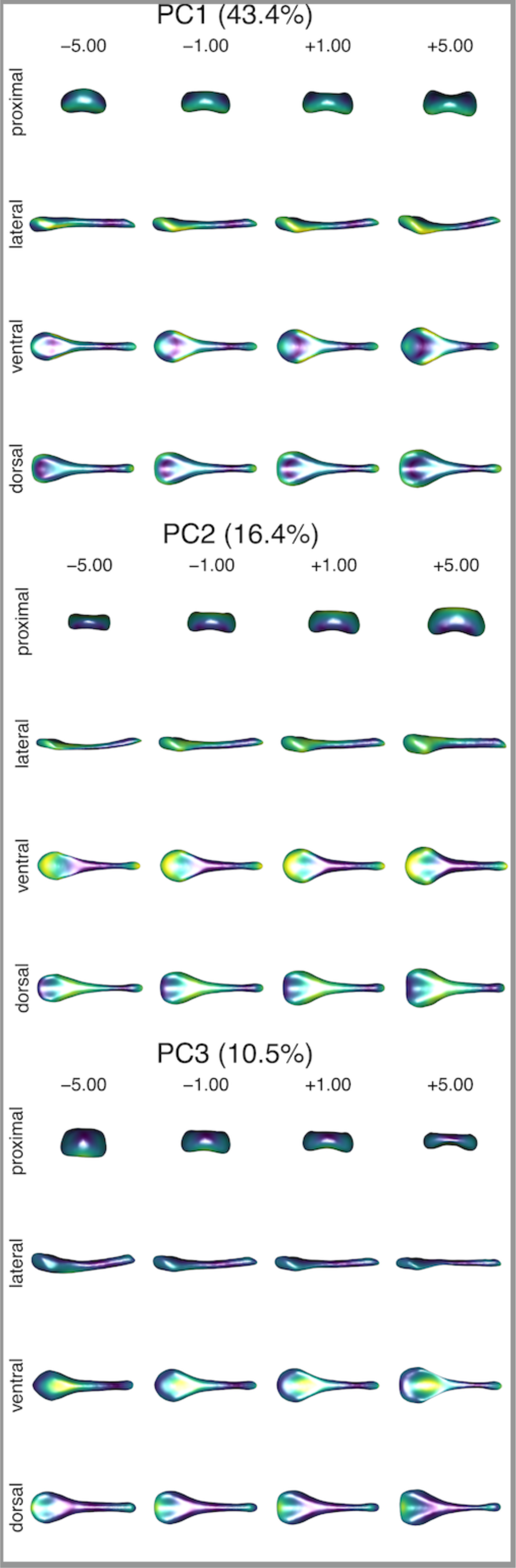
Variance map of bacula alignments. This figure is similar to figure 6, but these are projections of shape described by PC1, PC2, and PC3.

**Supplementary File 2.**
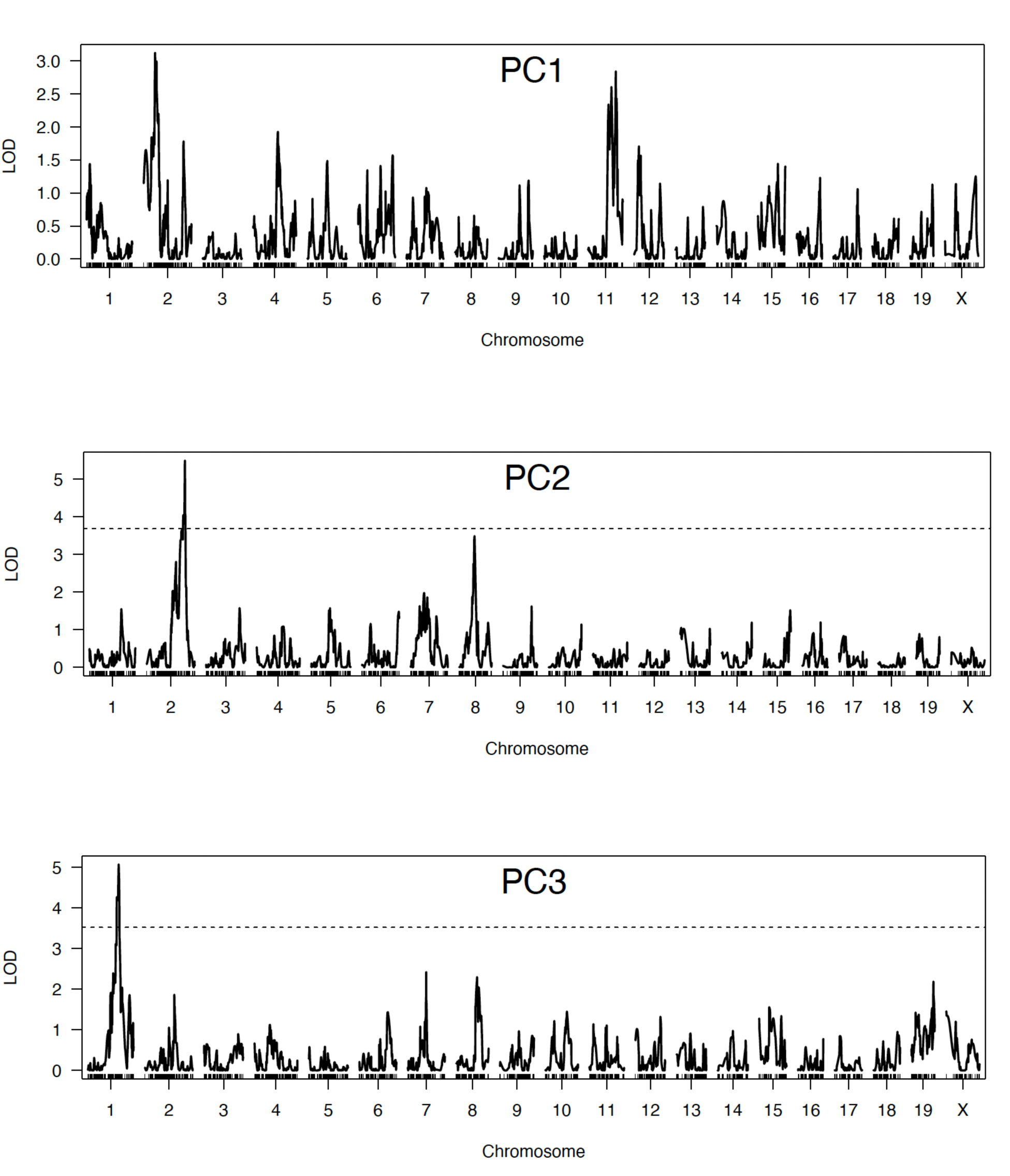
QTL map of baculum shape, described by PC1, PC2, and PC3.

## APPENDIX

### A1: Space partitioning tree

Given a pair of surfaces, naively searching for the best match for every point on each surface is an O(*n*^2^) operation, untenable for the large number of points (often in the hundreds of thousands to tens of millions) per surface. To bring the computational cost of finding the best match down to O(*n·*log *n*), we can use a hierarchal space partitioning tree – a data structure that repeatedly splits the data space and stores each data point appropriately in the resulting bins. After the hierarchal splitting is complete, we are left with a tree structure that can be traversed based on similarity measures (or other criteria) to find the terminal node containing the data most relevant to a query. Searching with this tree structure brings the complexity down to O(*n·*log *n*), much more acceptable for the scale of the data sets we are working with [41]. There are numerous algorithms for hierarchal space partitioning, but for our purposes, implementing a modified k-d tree is adequate. In a typical k-d tree, space is divided up by finding the median point on each dimensional axis, one at a time, in order to provide a balanced structure for optimal traversal time. k-d trees are not constrained to two or three dimensions; they are theoretically able to handle the addition of local shape descriptors. Normally, dimensions are split in turn, and it is also possible to allow overlapping nodes akin to those in an R-tree to partition for more robust results.

Choosing the optimal axis to split the data is not a trivial problem in the context of GPSA2, where dimensionality can grow significantly with the addition of landmark-based descriptors. Even after standardizing the dimensions (dividing by the standard deviation, later in these appendices this is referred to as “standardized space”), a normal k-d tree implementation faced stability and balancing issues. Our solution was to define a hyperplane perpendicular to the longest axis in standardized space for the data within the node (found using Eigen decomposition) containing the centroid of the data in the node. Checking which side of the hyperplane a given data point lies on is a simple dot product operation between the vector normal to the hyperplane and the difference between the point and the node centroid, all scaled using the same standard deviations as above. This method proved to be much more stable and similarly efficient in construction and traversal for a low number of descriptors but grew in computational cost with a large number of descriptors due to the decomposition step. To enable virtually overlapping nodes during recursive traversal (overlapping the nodes in the data structure via record duplication be prohibitively memory intensive), we define a scalar parameter r that is multiplied by the standardized distance between the centroids of the two child nodes of the current node to define a buffer zone where both halves of a node are searched, rather than only one side. The results are compared, and the best match is passed back up to the parent. The size of this buffer zone must be weighed carefully against the number of points represented by the tree to avoid very long run times. Scans with tens of thousands of points supported an overlap of value of r = 1.0, while scans with millions of points were only computationally bearable with an overlap value of r = 0.1.

Once the best match for a given point has been found using the tree, the given point is projected onto each of the faces adjoining the matching point. The projected point with the shortest distance in Euclidean space is used as the final match. The barycentric coordinates of this projection are used to interpolate pointwise properties for later use. Euclidean distance is used rather than the standardized difference because finding the optimal projection for the interpolated standardized difference over a triangle is a quadratic programming problem that is computationally expensive to solve for every point. This approximation is acceptable because highly curved or detailed regions tend to require denser sampling (i.e., smaller faces) for an accurate representation, meaning the error in the projection is constrained by the relatively smaller size of the face. In flatter, more homogenous regions that can be represented by fewer, larger faces, the reverse is true: a projection is necessary to minimize the error in Euclidean space, which would be the source of most of the error in a standardized distance measure. This constraint on error is only logical for geometry-based local shape descriptors; for landmark-based descriptors, subdividing large faces before analysis could help constrain such error. In future implementations, virtual subdivisions of such faces could skirt the quadratic programming problem while enforcing an upper bound on the potential per-point error in homology approximation.

### A2: Match quality metric

For the purpose of resistant-fit superimposition between two surfaces, we must be able to compare one pair of matched points in an approximated homology map to another pair so that our algorithm can determine which pairs to aggregate into the superimposition cost function and which should be ignored. For a given point *q*, we not only want to know how far away it is in standardized space from its best match, *qʹ*, we also want to know what the best match for *qʹ*, *qʹʹ*, is back on the surface containing *q*, and how far away *qʹʹ* is from *q* (Fig. A1).

**Figure A1:**
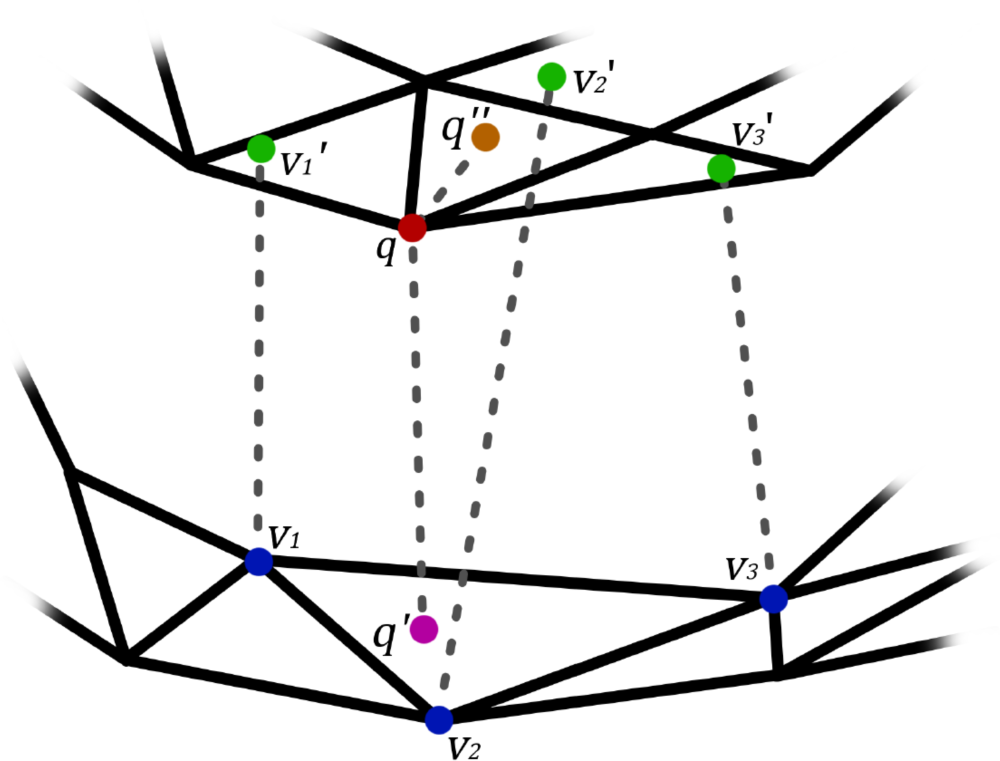
A diagram depicting the original point *q* (red), its best match *qʹ* (purple), the vertices of the face the best match lies on, *v_1_*, *v_2_*, and *v_3_* (blue), the best matches for those vertices, *v_1_ʹ*, *v_2_ʹ*, and *v_3_ʹ* (green), and the interpolation of those matches, *qʹʹ* (orange). The match quality metric is a function of the distance between *q* and *qʹ* and the distance between *q* and *qʹʹ*.

Space partitioning trees are constructed for each surface, so we could theoretically run *qʹ* through the space partitioning tree for the surface containing *q*, but a computationally cheaper alternative is to find the best matches for every point on each surface (which we need to do anyway for superimposition and shape distance measurement), store said matches, then use the barycentric coordinates {*b_1_*, *b_2_*, *b_3_*} from projecting *qʹ* onto the face *f* containing *qʹ* as weights for a linear interpolation of the matches for the vertices of face *f*. We’ll label these vertices *v_1_*, *v_2_*, and *v_3_*, and the best matches for these vertices *v_1_ʹ*, *v_2_ʹ*, and *v_3_ʹ*, respectively. The equation for this interpolated best match, *qʹʹ* is simply:

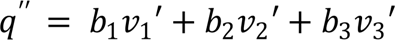

We may reasonably define *qʹ* as “good match” for point *q* if it is close to *q* in standardized space: this means that the two points are similar in terms of position in 3D space and in terms of shape descriptors as weighted by our chosen parameterization. However, given that *qʹ* is the best match we can achieve for *q* using our space partitioning method, it is also reasonable to state that even if *qʹ* is not very close to *q*, they may still be the best match possible, merely separated by distance. This can occur when homologous regions have some local shape variation that causes them to be locally misaligned in the optimal global superimposition. Recognizing this, we can also reasonably state that if *qʹʹ* is close to *q* in standardized space, meaning the matches are in close agreement with one another, then the pair can still be considered a good match. We can already measure the standardized distance *D_s_* (main text) between two points. Mathematically, then, if either or both of *D_s_*(*q*, *qʹ*) or *D_s_*(*q*, *qʹʹ*) is small, we want our metric to be small as well. A simple function that satisfies this requirement is the product of these two standardized distances. In our resulting match quality metric, we also take the square root in order to maintain keep the values well-behaved for the rest of the resistant-fit algorithm when both *D_s_*(*q*, *qʹ*) and *D_s_*(*q*, *qʹʹ*) are not small:

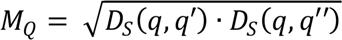

## Notes

### Competing Interest Statement

The authors have declared no competing interest.

